# Viral and bacterial infection elicit distinct changes in plasma lipids in febrile children

**DOI:** 10.1101/655704

**Authors:** Xinzhu Wang, Ruud Nijman, Stephane Camuzeaux, Caroline Sands, Heather Jackson, Myrsini Kaforou, Marieke Emonts, Jethro Herberg, Ian Maconochie, Enitan D Carrol, Stephane C Paulus, Werner Zenz, Michiel Van der Flier, Ronald de Groot, Federico Martinon-Torres, Luregn J Schlapbach, Andrew J Pollard, Colin Fink, Taco T Kuijpers, Suzanne Anderson, Matthew Lewis, Michael Levin, Myra McClure, on behalf of EUCLIDS consortium

**Affiliations:** Jefferiss Research Trust Laboratories, Department of Medicine, Imperial College London; Section of Paediatrics, Department of Medicine, Imperial College London; National Phenome Centre and Imperial Clinical Phenotyping Centre, Department of Surgery and Cancer, IRDB Building, Du Cane Road, Imperial College London, London, W12 0NN, United Kingdom; Great North Children’s Hospital, Paediatric Immunology, Infectious Diseases & Allergy, Newcastle upon Tyne Hospitals NHS Foundation Trust, Newcastle upon Tyne, United Kingdom; Institute of Cellular Medicine, Newcastle University, Newcastle upon Tyne, United Kingdom; NIHR Newcastle Biomedical Research Centre based at Newcastle upon Tyne Hospitals NHS Trust and Newcastle University, Newcastle upon Tyne, United Kingdom; Department of Paediatric Emergency Medicine, St Mary’s Hospital, Imperial College NHS Healthcare Trust, London, United Kingdom; Institute of Infection and Global Health, University of Liverpool, Liverpool, United Kingdom; Department of Infectious Diseases, Alder Hey Children’s NHS Foundation Trust, Liverpool, United Kingdom; Liverpool Health Partners, Liverpool, United Kingdom; Department of General Paediatrics, University of Graz, Graz, Austria; Pediatric Infectious Diseases and Immunology, Wilhelmina Children’s Hospital, University Medical Center Utrecht, Utrecht, The Netherlands; Pediatric Infectious Diseases and Immunology, Amalia Children’s Hospital, and Section Pediatric Infectious Diseases, Laboratory of Medical Immunology, Department of Laboratory Medicine, Radboud Institute for Molecular Life Sciences, Radboud University Medical Center, Nijmegen, The Netherlands; Pediatrics Department, Translational Pediatrics and Infectious Diseases Section, Santiago de Compostela, Spain; Faculty of Medicine, University of Queensland, Brisbane, QLD, Australia; Department of Paediatrics, University of Oxford and the NIHR Oxford Biomedical Research Centre, Oxford, United Kingdom; Micropathology Ltd, University of Warwick, Warwick, United Kingdom; Division of Pediatric Hematology, Immunology and Infectious diseases, Emma Children’s Hospital Academic Medical Center, Amsterdam, The Netherlands; Medical Research Council Unit Gambia, Banjul, The Gambia

## Abstract

Fever is the most common reason that children present to Emergency Departments in the UK. Clinical signs and symptoms suggestive of bacterial infection are often non-specific, and there is no definitive test for the accurate diagnosis of infection. As a result, many children are prescribed antibiotics often unnecessarily, while others with life-threatening bacterial infections can remain untreated. The ‘omics’ approaches to identifying biomarkers from the host-response to bacterial infection are promising. In this study, lipidomic analysis was carried out with plasma samples obtained from febrile children with confirmed bacterial infection (n=20) and confirmed viral infection (n=20). We show for the first time that bacterial and viral infection elicit distinct changes in the host lipidome. Glycerophosphoinositol, sphingomyelin, lysophosphotidylcholine and cholesterol sulfate were increased in the confirmed virus infected group, while fatty acids, glycerophosphocholine, glycerophosphoserine, lactosylceramide and bilirubin were increased in cases with confirmed bacterial infection. A combination of three lipids achieved the area under the receiver operating characteristic (ROC) curve of 0.918 (95% CI 0.835 to 1). This pilot study demonstrates the potential of metabolic biomarkers to assist clinicians in distinguishing bacterial from viral infection in febrile children, to facilitate effective clinical management and to the limit inappropriate use of antibiotics.

## Introduction

Fever is the one of the most common reasons that children present to Emergency departments in hospitals, especially in children under 5 years of age, in England [1] and in the US [2]. Serious bacterial infection accounts for 5-15% of the febrile children presenting [3][4][5] and most cases originating from a viral aetiology are self-limiting. Currently bacterial infection is confirmed by positive microbiological culture of a sterile sample (blood, clean catch urine or cerebrospinal fluids (CSF)). However, this can take 24-48 hours and is compounded by having a high false-negative [4,6] and false positive [7] rates by contaminating pathogens. Molecular detection of specific pathogens is an option but results can be confounded by co-infections and samples need to be obtained from the site of infection which can be both invasive and impractical [8]. Because it is challenging for paediatricians to differentiate between bacterial and viral infection in acute illness, antibiotics are often prescribed as a precautionary measure, contributing to the rise of antimicrobial resistance.

It is clear that reliable biomarkers are urgently needed that distinguish bacterial from viral infection for the purpose of good clinical management and reducing antibiotic use. Host biomarkers, i.e. the physiological changes of the host in response to a specific pathogen, have untapped diagnostic potential and their discovery can be accelerated by the advances in ‘omics’ research, especially in the field of transcriptomics [9–12] and proteomics [13–15]. Metabolomics has the added advantage that it is considered to most closely reflect the native phenotype and functional state of a biological system. One *In vivo* animal study revealed that distinct metabolic profiles can be derived from mice infected with different bacteria [16] and several similar studies focusing on meningitis have shown that metabolic profiling of CSF can differentiate between meningitis and negative controls [17], as well as between viral and bacterial meningitis [18]. Mason *et al* [19] demonstrated the possibility of diagnosis and prognosis of tuberculous meningitis with non-invasive urinary metabolic profiles. Metabolic changes in urine can be used to differentiate children with respiratory syncytial virus (RSV) from healthy control, as well as from those with bacterial causes of respiratory distress [20].

Lipids are essential structural components of cell membranes and energy storage molecules. Thanks to the advances in lipidomics, a subset of metabolomics, lipids and lipid mediators have been increasingly recognised to play a crucial role in different metabolic pathways and cellular functions, particularly in immunity and inflammation [21,22]. However, the potential of lipidomics to distinguish bacterial from viral infection in febrile children has never been explored.

In this study, we undertook a lipidomic analysis of plasma taken from febrile children with confirmed bacterial infection (n=20) and confirmed viral infection (n=20) as a proof of concept study. We show that bacterial and viral infection elicit distinct changes in the plasma lipids of febrile children that might be exploited diagnostically.

## Methods

### Study population and sampling

The European Union Childhood Life-Threatening Infectious Disease Study (EUCLIDS) [23] prospectively recruited patients, aged from 1 month to 18 years, with sepsis or severe focal infection from 98 participating hospitals in the UK, Austria, Germany, Lithuania, Spain and the Netherlands between 2012 and 2015. Plasma and other biosamples were collected to investigate the underlying genetics, proteomics and metabolomics of children with severe infectious disease phenotype.

Infections in Children in the Emergency Department (ICED) study aimed to define clinical features that would predict bacterial illness in children and patterns of proteomics, genomics and metabolomics associated with infections. This study included children aged 0 −16 years at Imperial College NHS Healthcare Trust, St Mary’s Hospital, between June 2014 and March 2015 [24].

The population consisted of children (≤17 years old) presenting with fever ≥ 38 °C, with diverse clinical symptoms and a spectrum of pathogens. Both studies were approved by the local institutional review boards (ICED REC No 14/LO/0266; EUCLIDS REC No 11/LO/1982). Written informed consent was obtained from parents and assent from children, where appropriate. For the EUCLIDS study, a common clinical protocol agreed by EUCLIDS Clinical Network and approved by the Ethics Committee was implemented at all hospitals.

Patients were divided into those with confirmed bacterial (n=20) and confirmed viral (n=20) infection groups. The bacterial group consisted exclusively of patients with confirmed sterile site culture-positive bacterial infections, and the viral infection group consisted of only patients with culture, molecular or immunofluorescent-confirmed viral infection and having no co-existing bacterial infection.

Blood samples were collected in tubes spray-coated with EDTA at, or as close as possible to, the time of presentation to hospital and plasma obtained by centrifugation of blood samples for 10 mins at 1,300 g at 4 °C. Plasma was stored at − 80°C before being shipped on dry ice to Imperial College London for lipidomic analysis.

### Lipidomic analysis

Lipidomic analysis was carried out as previously described [25]. Briefly, 50 μl of water were added to 50 μl of plasma, vortexed and shaken for 5 min at 1,400 rpm at 4°C. Four hundred μl of isopropanol containing internal standards (9 in negative mode, 11 for positive mode covering 10 lipid sub-classes) were added for lipid extraction. Samples were shaken at 1,400 rpm for 2 hours at 4°C then centrifuged at 3,800 g for 10 min. Two aliquots of 100 μl of the supernatant fluid were transferred to a 96-well plate for ultra-performance liquid chromatography (UPLC) −mass spectrometry (MS) lipidomics analysis in positive and negative mode.

Liquid chromatography separation was carried out using an Acquity UPLC system (Waters Corporation, USA) with an injection volume of 1μl and 2μl for Positive and Negative ESI, respectively. An Acquity UPLC BEH column (C8, 2.1 × 100 mm, 1.7 μm; Waters Corporation, USA) was used for the purpose. Mobile phase A consisted of water/isopropanol/acetonitrile (2:1:1; v:v:v) with the addition of 5 mM ammonium acetate, 0.05% acetic acid and 20 μM phosphoric acid. Mobile phase B consisted of isopropanol: acetonitrile (1:1; v:v) with the addition of 5mM ammonium acetate and 0.05% acetic acid. Flow rate was 0.6 ml/min with a total run time of 15 min and the gradient set as starting condition of 1% mobile phase B for 0.1 min, followed by an increase to 30% mobile phase B from 0.1 to 2 min, and to 90% mobile phase B from 2 min to 11.5 min. The gradient was held at 99.99% mobile phase B between 12 and 12.55 min before returning to the initial condition for re-equilibrium.

MS detection was achieved using a Xevo G2-S QTof mass spectrometer (Waters MS Technologies, UK) and data acquired in both positive and negative modes. The MS setting was configured as follows: capillary voltage 2.0 kV for Positive mode, 1.5 kV for Negative mode, sample cone voltage 25V, source offset 80, source temperature 120 °C, desolvation temperature 600 °C, desolvation gas flow 1000 L/h, and cone gas flow 150 L/h. Data were collected in centroid mode with a scan range of 50 −2000 m/z and a scan time of 0.1s. LockSpray mass correction was applied for mass accuracy using a 600 pg/ μL leucine enkephaline (m/z 556.2771 in ESI+, m/z 554.2615 in ESI-) solution in water/acetonitrile solution (1:1; v/v) at a flow rate of 15 μl/min.

### Spectral and statistical analysis

A Study Quality Control sample (SQC) was prepared by pooling 25 μl of all samples. The SQC was diluted to seven different concentrations, extracted at the same ratio 1:4 with isopropanol and replicates acquired at each concentration at the beginning and end of the run. A Long-Term Reference sample (LTR, made up of pooled plasma samples from external sources) and the SQC were diluted with water (1:1; v:v) and 400 μL of isopropanol containing internal standards (the same preparation as for the study samples) and injected once every 10 study samples, with 5 samples between a LTR and a SQC. Deconvolution of the spectra was carried out using the XCMS package. Extracted metabolic features were subsequently filtered and only those present with a relative coefficient of variation less than 15% across all SQC samples were retained. Additionally, metabolic features that did not correlate with a coefficient greater than 0.9 in a serial dilution series of SQC samples were removed.

Multivariate data analysis was carried out using SIMCA-P 14.1 (Umetrics AB, Sweden). The dataset was pareto-scaled prior to principal component analysis (PCA) and orthogonal partial least squares discriminate analysis (OPLS-DA). While PCA is an unsupervised technique useful for observing inherent clustering and identifying potential outliers in the dataset, OPLS-DA is a supervised method in which data is modelled against a specific descriptor of interest (in this case viral vs. bacterial infection classes). As for all supervised methods, model validity and robustness must be assessed before results can be interpreted. For OPLS-DA, model quality was assessed by internal cross-validation (Q^2^Y-hat value) and permutation testing in which the true Q^2^Y-hat value is compared to 999 models with random permutations of class membership. For valid and robust models (positive Q^2^Y-hat and permutation p-value < 0.05), metabolic features responsible for class separation were identified by examining the corresponding S-plot (a scatter plot of model loadings and correlation to class) with a cut-off of 0.05.

### Metabolite annotation

Short-listed metabolic features were subjected to tandem mass spectrometry in order to obtain fragmentation patterns. Patterns were compared against metabolome databases (Lipidmaps, HMDB, Metlin). Isotopic distribution matching was also checked. In addition, when possible the fragmented patterns were matched against available authentic standards run under the same analytical setting for retention time and MS/MS patterns. Annotation level, according to the Metabolomics Standards Initiative, are summarised in Table 2 [26].

### Single feature ROC curve analysis

Analysis was performed with the web server, MetaboAnalyst 4.0. Sensitivities and specificities of lipids and predicted probabilities for the correct classification were presented as Receiver Operating Characteristic (ROC) curves. The Area Under the Curve (AUC) represents the discriminatory power of the lipids, with the value closest to 1 indicating the better classification.

### Feature Selection

In order to identify a small diagnostic signature capable of differentiating between viral and bacterial infections, an ‘in-house’ variable selection method, forward selection-partial least squares (FS-PLS; https://github.com/lachlancoin/fspls.git), was used that eliminates highly correlated features. The first iteration of FS-PLS considers the levels of all features (N) and initially fits N univariate regression models. The regression coefficient for each model is estimated using the Maximum Likelihood Estimation (MLE) function, and the goodness of fit is assessed by a t-test. The variable with the highest MLE and smallest p-value is selected first (SV1). Before selecting which of the N-1 remaining variables to use next, the algorithm projects the variation explained by SV1 using Singular Value Decomposition. The algorithm iteratively fits up to N-1 models, at each step projecting the variation corresponding to the already selected variables, and selecting new variables based on the residual variation. This process terminates when the MLE p-value exceeds a pre-defined threshold (p_thresh_). The final model includes regression coefficients for all selected variables. The sensitivity and specificity of the lipid signature identified by FS-PLS were presented as a receiver operating characteristic (ROC) curve.

## Results

### Patient characteristics

The baseline characteristics were divided into those with definitive bacterial and definitive viral infection, summarised in Table 1. When selecting patient samples, patient characteristics were matched as much as possible to ensure no particular factor would confound the model. There was no significant difference in ages between the two groups (p=0.97). Both groups had similar gender split. Seven from definitive bacterial infection group and 6 from the definitive viral infection group were admitted to the Paediatric Intensive Care Unit (PICU). A range of pathogens was present in each group.

**Table 1.**
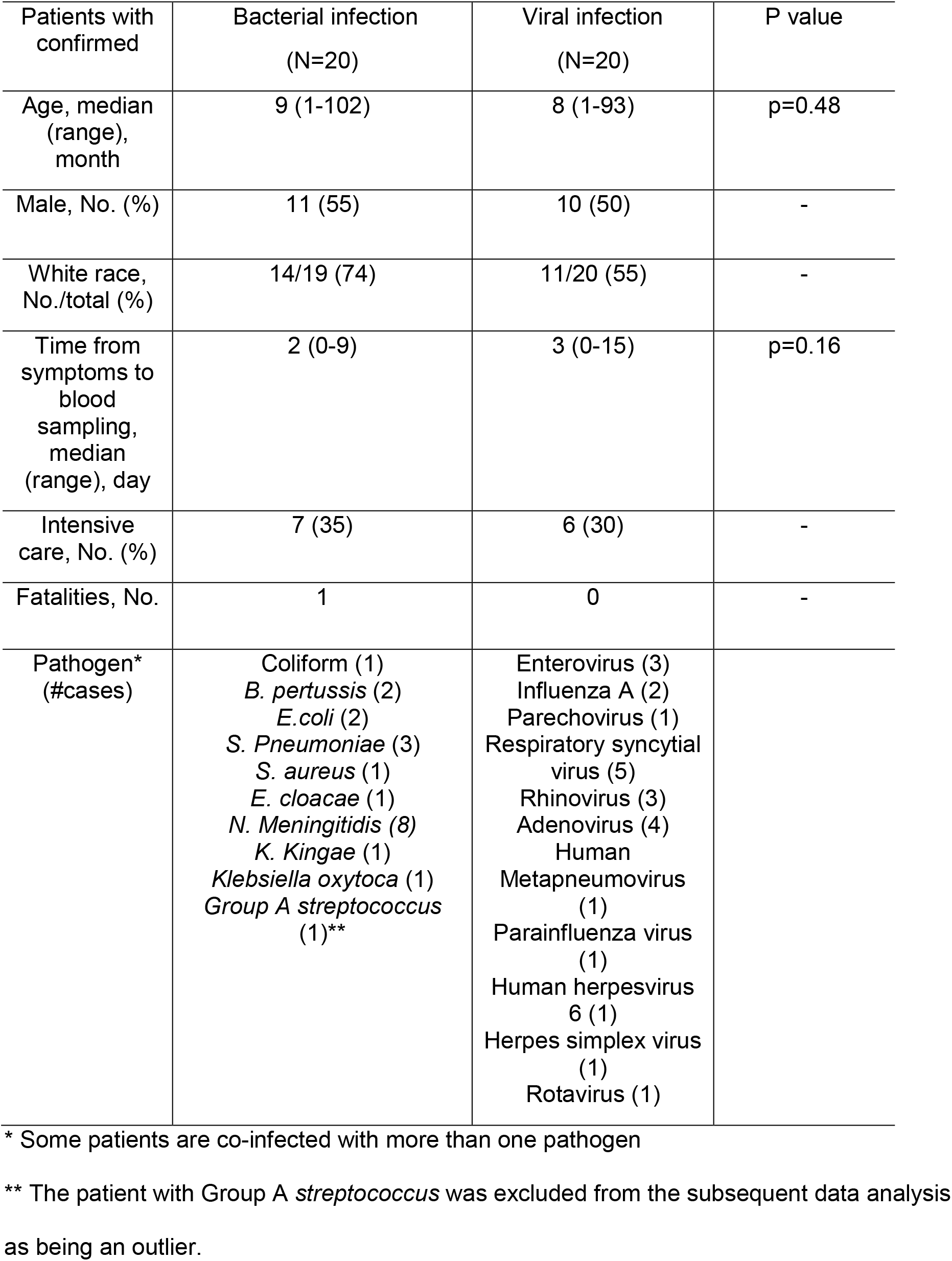
Demographic and clinical patient characteristic.

### Plasma Lipidome can differentiate bacterial from viral infection

PCA was conducted first to evaluate the data, visualise dominant patterns, and identify outliers within populations (Figure 1). The same outlier sample was present in both negative (Figure 1A) and positive (Figure 1B) polarity datasets and as such, was removed from subsequent analysis. SQC samples were tightly grouped together in the PCA scatter plot, indicating minimum analytical variability throughout the run.

**Figure 1.**
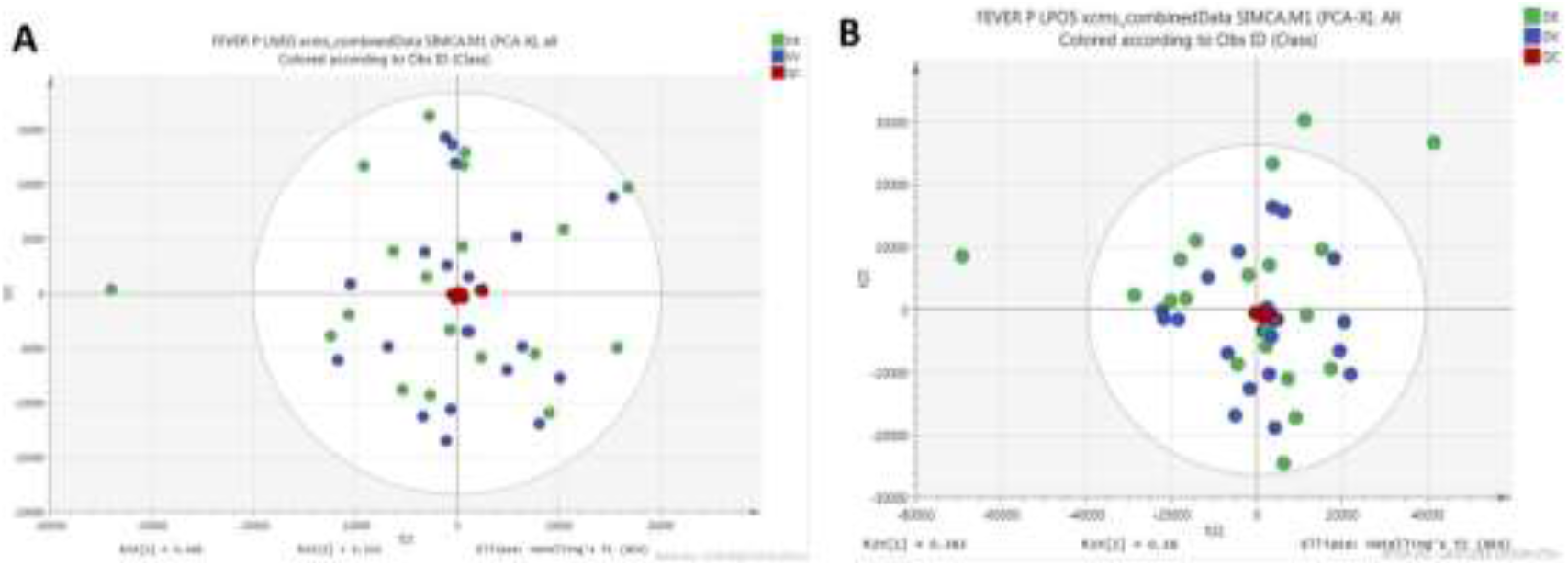
Principal components analysis (PCA) of lipidomics dataset. (A) Scatter plot of PCA model from data acquired in negative polarity mode. (B) Scatter plot of PCA model from data acquired in positive polarity mode. Quality control samples are shown in red, bacterial infected samples are shown in blue and viral infected samples shown in green.

OPLS-DA, a supervised PCA method, was carried out on both positive and negative polarity datasets. In the positive polarity mode no model was successfully built to distinguish between viral and bacterial infection groups (data not shown). However, in the negative polarity dataset, an OPLS-DA model separated bacterial infected samples from viral infected samples. The robustness of the model was characterised by R2X (cum) = 0.565, R2Y-hat (cum)= 0.843 and Q2Y-hat (cum)= 0.412 and permutation p-value=0.01 (999 tests). Cross-validated scores plot using the whole lipidome dataset indicated bacterial infected samples were more prone to miss-classification than viral infected samples (Figure 2A).

**Figure 2.**
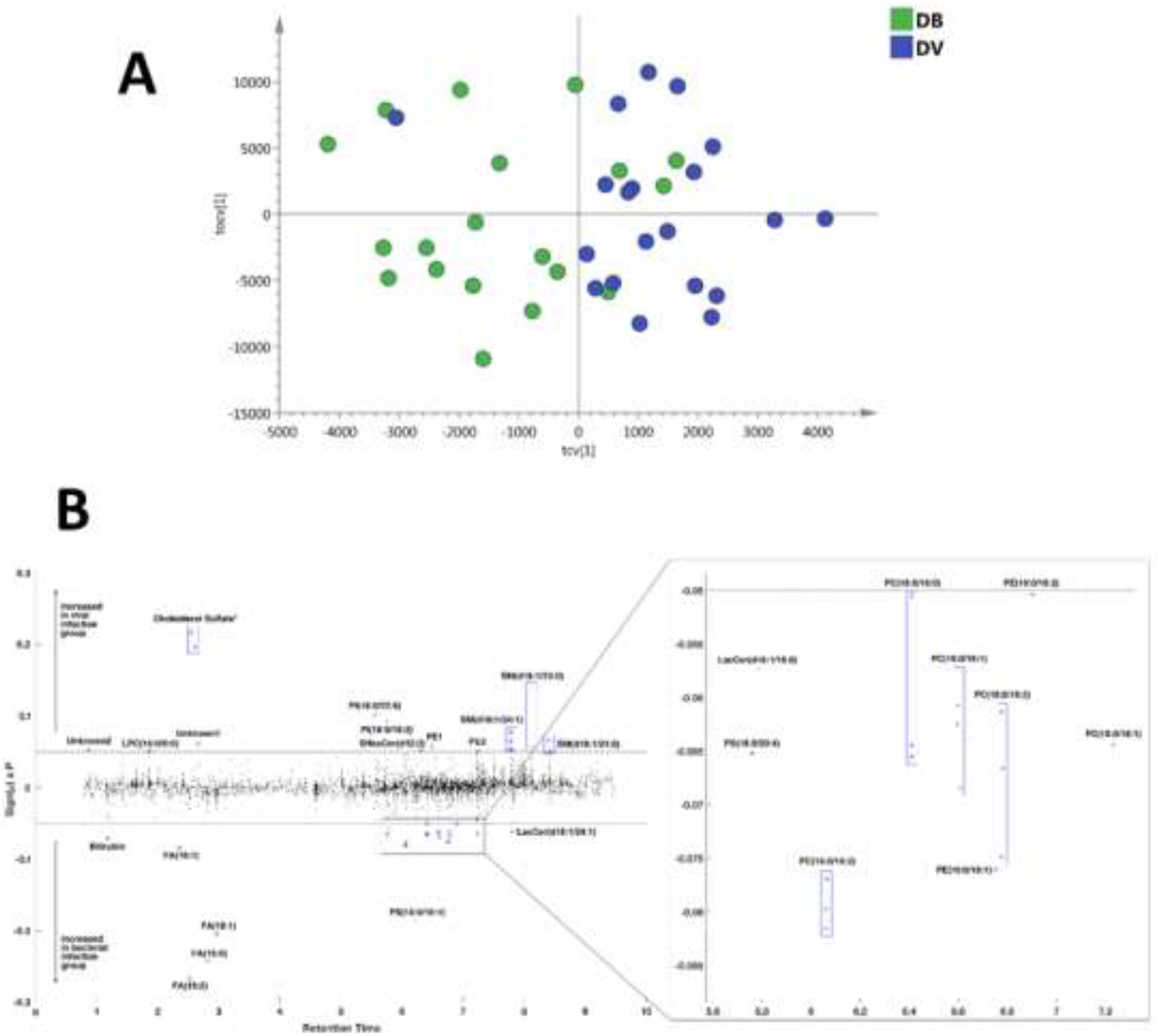
Supervised pattern recognition results of negative polarity lipidomics dataset. (A) The scatter plot of the cross-validated score vectors showing the clustering of definitive bacterial infected samples (green dots) from definitive viral infected samples (blue dots). (B) Manhattan-style plot of the 3891 lipid features detected by lipid-positive mode UPLC-MS with 40 features showing a significant association with infection type (as determined by model S-plot) highlighted and annotated. *Cholesterol sulfate – isomers due to different position of the sulfate.

### Lipid changes were not the same in the bacterial and viral infected groups

Metabolic features contributing to the separation of the model are plotted in Figure 2B and summarised in Table 2. Glycerophosphoinositol, monoacylglycerophosphocholine, sphingomyelin and sulfatide were only increased in the viral group, while fatty acids, glycerophosphocholine, glycerophosphoserine and lactosylceramide were only increased in bacterial infection. Bilirubin and cholesterol sulfate, although not lipids, were detected by lipidomic analysis, and these were increased in the bacterial and viral groups, respectively.

**Table 2.** Metabolic features changed in bacterial and viral group.

|  | m/z | retention time | annotation | annotation level | ion type | neutral formula |
| --- | --- | --- | --- | --- | --- | --- |
| **Lipids/metabolites increased in bacterial infected group** | 279.231 | 2.52 | FA(18:2) | 2 | [M-H]- | C18H32O2 |
|  | 255.232 | 2.82 | FA(16:0) | 2 | [M-H]- | C16H32O2 |
|  | 281.247 | 2.96 | FA(18:1) | 2 | [M-H]- | C18H34O2 |
|  | 788.545 | 6.22 | PS(18:0/18:1) | 2 | [M-H]- | C39H74NO8P |
|  | 253.216 | 2.35 | FA(16:1) | 2 | [M-H]- | C16H30O2 |
|  | 742.54 | 6.06 | PC(16:0/18:2) | 2 | [M-CH3]- | C42H80NO8P |
|  | 716.524 | 6.75 | PE(16:0/18:1) | 2 | [M-H]- | C39H76NO8P |
|  | 583.256 | 1.18 | Bilirubin | 2 | [M-H]- | C33H36N4O6 |
|  | 810.53 | 5.76 | PS(18:0/20:4) | 2 | [M-H]- | C44H78NO10P |
|  | 846.624 | 7.23 | PC(18:0/18:1) | 2 | [M+PO4H2]- | C44H86NO8P |
|  | 1068.7 | 7.80 | LacCer(d18:1/24:1) | 2 | [M+PO4H2]- | C54H101NO13 |
|  | 770.571 | 6.78 | PC(18:0/18:2) | 2 | [M-CH3]- | C44H84NO8P |
|  | 744.556 | 6.60 | PC(16:0/18:1) | 2 | [M-CH3]- | C42H82NO8P |
|  | 958.589 | 5.78 | LacCer(d18:1/16:0) | 2 | [M+PO4H2]- | C46H87NO13 |
|  | 718.54 | 6.41 | PC(16:0/16:0) | 2 | [M-CH3]- | C40H80NO8P |
|  | 742.54 | 6.90 | PE(18:0/18:2) | 2 | [M-H]- | C41H78NO8P |
| **Lipids/metabolites increased in viral infected group** | 465.303 | 2.55 | Cholesterol sulfate | 2 | [M-H]- | C27H46O4S |
|  | 465.303 | 2.61 | Cholesterol sulfate | 2 | [M-H]- | C27H46O4S |
|  | 909.551 | 5.56 | PI(18:0/22:6) | 2 | [M-H]- | C49H83O13P |
|  | 861.55 | 5.75 | PI(18:0/18:2) | 2 | [M-H]- | C45H83O13P |
|  | 797.655 | 7.78 | SM(d18:1/24:1) | 2 | [M-CH3]- | C47H93N2O6P |
|  | 339.231 | 2.66 | UNKNOWN1 | 4 |  |  |
|  | 772.529 | 6.49 | PE1 | 3 | [M-H]- | C45H76NO7P |
|  | 897.648 | 8.12 | SM(d18:1/23:0) | 2 | [M+PO4H2]- | C46H93N2O6P |
|  | 239.157 | 0.87 | UNKNOWN2 | 4 |  |  |
|  | 886.609 | 6.31 | SHexCer(d42:3) | 2 | [M-H]- | C48H89NO11S |
|  | 554.346 | 1.86 | LPC(16:0/0:0) | 2 | [M+CH3COO]- | C24H50NO7P |
|  | 799.671 | 8.41 | SM(d18:1/24:0) | 2 | [M-CH3]- | C47H95N2O6P |
|  | 750.545 | 7.24 | PE2 | 3 | [M-H]- | C41H78NO7P |

FA: fatty acid; PE: glycerophosphotidylethanolamine; PC: glycerophosphocholine; PS:
glycerophosphoserine; LacCer: lactosylceramide; PI: glycerophosphoinositol; SM:
sphingomyelin; LPC: Lysophosphotidylcholine; SHexCer: Sulfatides.

### Evaluation of diagnostic potential of metabolic biomarkers

ROC curve analysis was performed to evaluate the diagnostic potential of these lipids in distinguishing bacterial from viral infection. Out of all discriminatory lipids, PC (16:0/16:0), unknown feature m/z 239.157 and PE (16:0/18:2) generated the highest AUCs of 0.774 (CI, 0.6-0.902), 0.721 (CI, 0.545- 0.871) and 0.705 (CI, 0.52 – 0.849), respectively (Figure 3).

**Figure 3.**
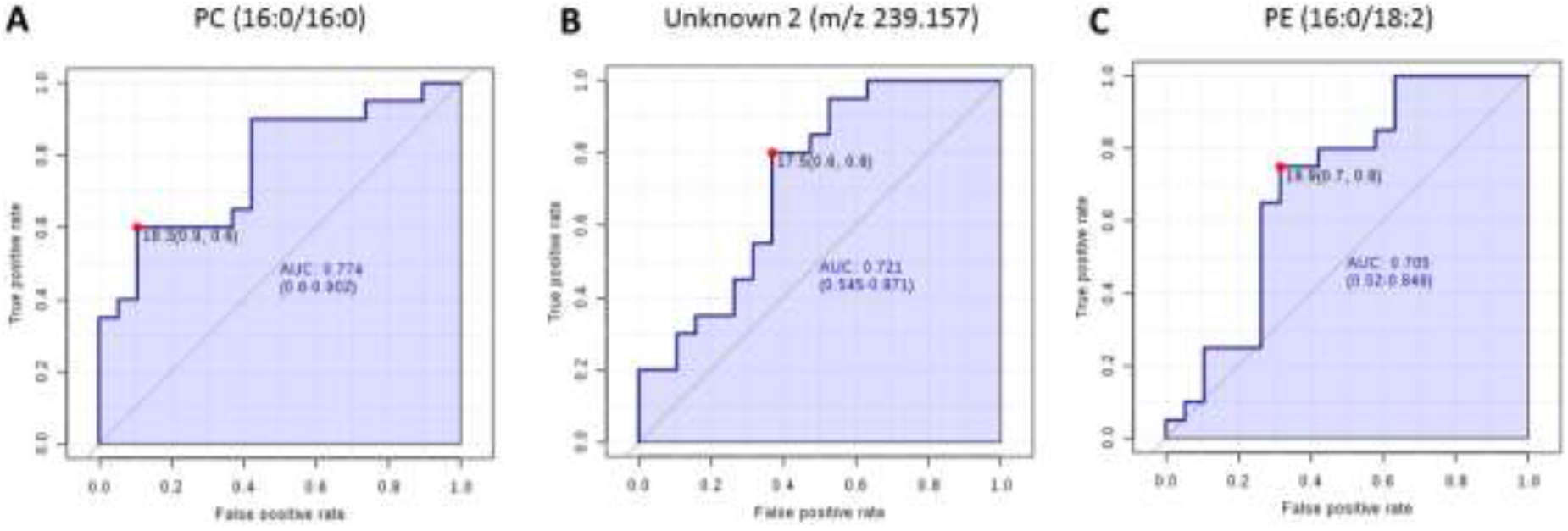
Receiver operator characteristic (ROC) analysis based on single lipids. ROC curve analysis of top 3 lipids PC (16:0/16:0) (A), unknown feature (m/z 239.157) (B) and PE (16:0/18:2) (C) which gave with highest Area Under the Curve (AUC) values.

FS-PLS was used to identify a small signature composed of non-correlated lipids that is capable of distinguishing between bacterial and viral samples. FS-PLS identified a signature made up the following 3 lipids: SHexCer(d42:3); PC (16:0/16:0); and LacCer(d18:1/24:1). This signature achieved an improved ROC curve with AUC of 0.9158 (95% confidence interval: 0.828 – 1) when compared with those generated from individual lipids (Figure 4).

**Figure 4.**
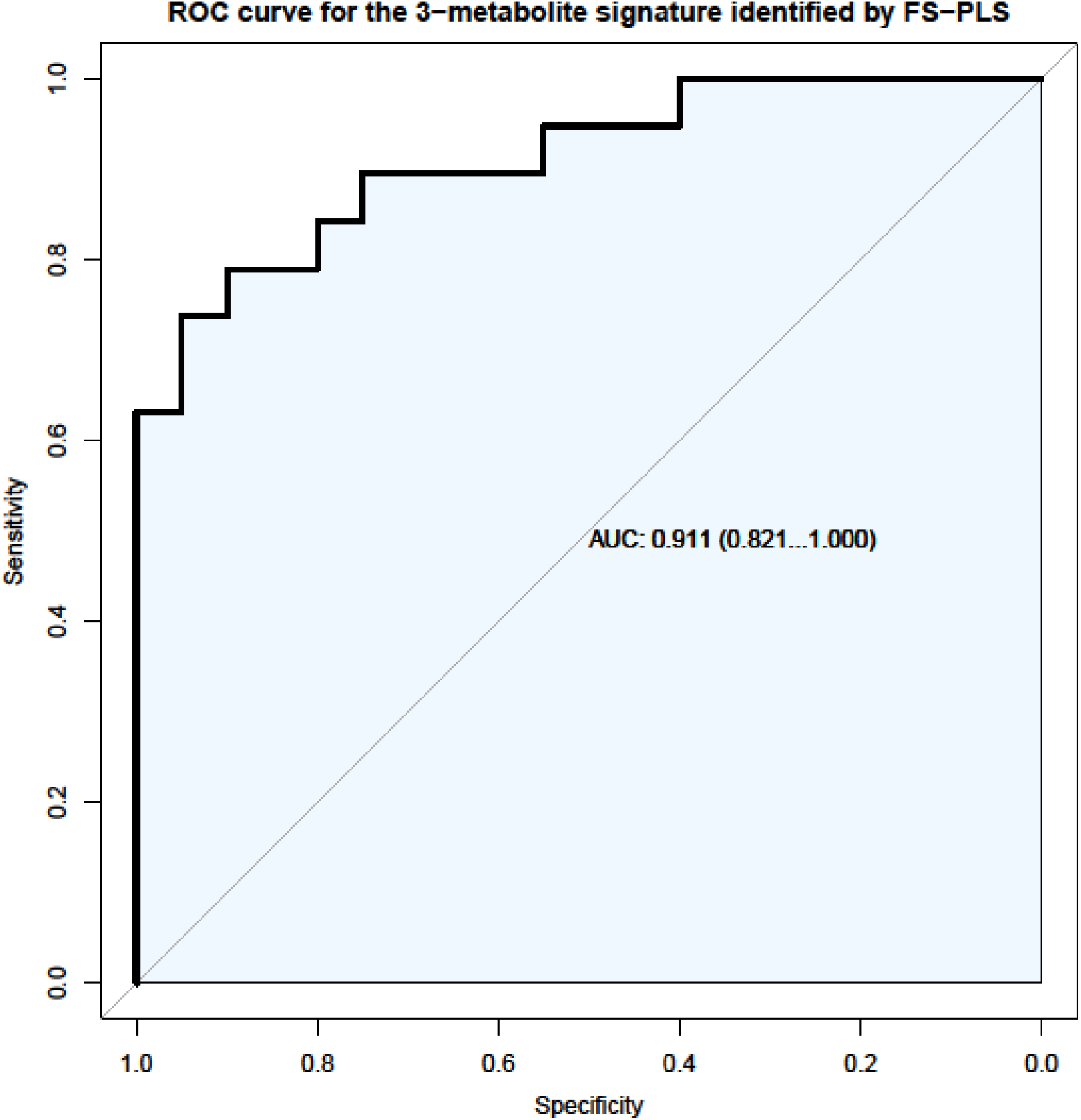
Receiver operator characteristic (ROC) analysis based on 3-lipid signature. A combination of SHexCer(d42:3), PC (16:0/16:0) and LacCer(d18:1/24:1) achieved AUC of 0.911 (CI 95% 0.821-1.000).

## Discussion

We have shown that differences in the host lipidome are induced by bacterial and viral infections. While differences in host responses between viral and bacterial infections have been previously reported, for example as differential expression of proteins, RNAs and level of metabolites [9–14,20], there have been no claims in relation to the lipidome changes in carefully-phenotyped samples. Although age is known to affect metabolism [27], it is important to note the metabolic changes associated with infection described herein, were consistent among samples from patients whose age ranged from 1 month to 9 years old.

Glycerophosphoinositol, sphingomyelin, lysophosphotidylcholine and cholesterol sulfate were increased in the confirmed virus-infected group, while fatty acids, glycerophosphocholine, glycerophosphoserine, lactosylceramide and bilirubin were increased in cases with confirmed bacterial infection.

The important effects of infection on fatty acid metabolism have been highlighted by Munger *et al* who demonstrated human cytomegalovirus (HCMV) up-regulated fatty acid biosynthesis in infected host cells. Pharmacologically inhibition of fatty acid biosynthesis suppressed viral replication for both HCMV and influenza A virus [28]. The importance of fatty acid biosynthesis may reflect its essential role in viral envelopment during viral replication. Rhinovirus induced metabolic reprogramming in host cell by increasing glucose uptake and indicated a shift towards lipogenesis and/or fatty acid update [29]. In our study, fatty acids linoleic acid (FA 18:2), palmitic acid (FA 16:0), oleic acid (FA 18:1) and palmitoleic acid (FA 16:1) were all decreased in viral infection (i.e. increased in bacterial infection), and may reflect enhanced lipogenesis and fatty acid uptake in the host cell during viral replication.

The increase in cholesterol sulfate observed may reflect changes in cellular lipid biosynthesis and T cell signalling during viral infection. Cholesterol sulfate is believed to play a key role as a membrane stabiliser [30] and can also act to modulate cellular lipid biosynthesis [31] and T cell receptor signal transduction [32]. Gong *et al* demonstrated that cholesterol sulfate was elevated in the serum of piglets infected with swine fever virus [33]. Taken together, these observations indicate that this compound could be a marker of viral infection.

The increase in sphingomyelin SM(d18:1/24:1), SM(d18:1/23:0) and SM(d18:1/24:0), and lysophosphocholine LPC (16:0) upon viral infection may also be linked to viral replication in infected cells. Accumulation of cone-shaped lipids, such as LPC in one leaflet of the membrane bilayer induces membrane curvature required for virus budding [34]. It is known that viral replication, for example in the case of dengue virus, induces dramatic changes in infected cells, including sphingomyelin, to alter the curvature and permeability of membranes [35]. Furthermore, the altered levels of sphingomyelin can be partially explained by elevated cytokine levels during bacterial infection, such as TNF-α [36], which can activate sphingomyelinase, hydrolysing sphingomyelin to ceramide [37]. Hence, sphingomyelin may be a class of lipids that plays a role in both viral and bacterial infection.

Lactosylceramide LacCer(d18:1/24:1) and LacCer (d18:1/16:0) were increased in bacterial infection. Lactosylceramide, found in microdomains on the plasma membrane of cells, is a glycosphingolipid consisting of a hydrophobic ceramide lipid and a hydrophilic sugar moiety. Lactosylceramide plays an important role in bacterial infection by serving as a pattern recognition receptors (PRRs) to detect pathogen-associated molecular patterns (PAMPs). Lactosylceramide composed of long chain fatty acid chain C24, such as LacCer(d:18:1/24:1) increased in our study, is essential for formation of LacCer-Lyn complexes on neutrophils, which function as signal transduction platforms for αMβ2 integrin-mediated phagocytosis [38].

Other lipids that were changed in our study, such as sulfatides and glycerophosphocolines, may also play an important role in bacterial infection. Sulfatides are multifunctional molecules involved in various biological process, including immune system regulation and during infection [39]. Sulfatides can act as glycolipid receptors that attach bacteria, such as *Escherichia coli* [40], *Mycoplasma hyopneumoniae* [41] and *Pseudomonas aeruginosa* [42] to the mucosal surfaces. Five glycerophosphocholine species including PC(16:0/18:2), PC(18:0/18:1), PC(18:0/18:2), PC(16:0/16:0) and PC(16:0/18:1) were increased in bacterial infected samples. The increase in glycerophosphocholine was demonstrated in a lipidomics study looking at plasma from tuberculosis patients [43], however, the exact role of glycerophosphocholine remains elusive. Bilirubin is detected as a consequence of breadth of lipidome coverage, and its role in infection is unclear. The lipid species identified in this study present an opportunity for further mechanistic study to understand the host responses in bacterial or viral infection.

A combination of three lipids achieved a strong area under the receiver operating characteristic (ROC) curve of 0.918 (95% CI 0.835 to 1). Similar approaches have been taken using routine laboratory parameters and more recently gene expression where 2-gene transcripts achieved an ROC curve of 0.95 (95% CI 0.94 −1) [11].The relevance of our data is that they provide the potential for a rapid diagnostic test with which clinicians could distinguish bacterial from viral infection in febrile children.

The study has limitations. Firstly, we were unable to annotate 4 of the 29 discriminatory features, of which two were assigned with only a broad lipid class by identifying the head group (PE). The unknown feature with m/z of 239.157 achieved the second highest AUC for ROC curve analysis on an individual basis. The unknown identity prevents this feature from being a potential marker and hinders biological understanding. This feature, however, was not included in the final 3-lipid panel that gave the highest AUC. Secondly, the sample size in this pilot study is small. Validation studies using quantitative assay are now required to confirm the findings.

This is the first lipidomics study carried out on plasma taken from febrile children for the purpose of distinguishing bacterial from viral infection. It demonstrates the potential of this approach to facilitate effective clinical management by rapidly diagnosing bacterial infection in paediatrics.

## Appendix: EUCLIDS Consortium members

EUCLIDS consortium (https://www.euclids-project.eu) is composed by:

### Imperial College partner (UK)

#### Members of the EUCLIDS Consortium at Imperial College London (UK) Principal and co-investigators

Michael Levin (grant application, EUCLIDS Coordinator)

Lachlan Coin (bioinformatics)

Stuart Gormley (clinical coordination)

Shea Hamilton (proteomics)

Jethro Herberg (grant application, PI)

Bernardo Hourmat (project management)

Clive Hoggart (statistical genomics)

Myrsini Kaforou (bioinformatics)

Vanessa Sancho-Shimizu (genetics)

Victoria Wright (grant application, scientific coordination)

### Consortium members at Imperial College

Amina Abdulla

Paul Agapow

Maeve Bartlett

Evangelos Bellos

Hariklia Eleftherohorinou

Rachel Galassini

David Inwald

Meg Mashbat

Stefanie Menikou

Sobia Mustafa

Simon Nadel

Rahmeen Rahman

Clare Thakker

## EUCLIDS UK Clinical Network

Poole Hospital NHS Foundation Trust, Poole: Dr S Bokhandi (PI), Sue Power, Heather Barham

Cambridge University Hospitals NHS Trust, Cambridge: Dr N Pathan (PI), Jenna Ridout, Deborah White, Sarah Thurston

University Hospital Southampton, Southampton: Prof S Faust (PI), Dr S Patel (coinvestigator), Jenni McCorkell.

Nottingham University Hospital NHS Trust: Dr P Davies (PI), Lindsey Crate, Helen Navarra, Stephanie Carter 2

University Hospitals of Leicester NHS Trust, Leicester: Dr R Ramaiah (PI), Rekha Patel

Portsmouth Hospitals NHS Trust, London: Dr Catherine Tuffrey (PI), Andrew Gribbin, Sharon McCready

Great Ormond Street Hospital, London: Dr Mark Peters (PI), Katie Hardy, Fran Standing, Lauren O’Neill, Eugenia Abelake

King’s College Hospital NHS Foundation Trust, London; Dr Akash Deep (PI), Eniola Nsirim

Oxford University Hospitals NHS Foundation Trust, Oxford Prof A Pollard (PI), Louise Willis, Zoe Young

Kettering General Hospital NHS Foundation Trust, Kettering: Dr C Royad (PI), Sonia White Central Manchester NHS Trust, Manchester: Dr PM Fortune (PI), Phil Hudnott

### SERGAS Partner (Spain)

#### Principal Investigators

Federico Martinón-Torres1

Antonio Salas1,2

#### GENVIP RESEARCH GROUP (in alphabetical order)

Fernando Álvez González1, Ruth Barral-Arca1,2, Miriam Cebey-López1, María José Curras-Tuala1,2, Natalia García1, Luisa García Vicente1, Alberto Gómez-Carballa1,2, Jose Gómez Rial1, Andrea Grela Beiroa1, Antonio Justicia Grande1, Pilar Leboráns Iglesias1, Alba Elena Martínez Santos1, Federico Martinón-Torres1, Nazareth MartinónTorres1, José María Martinón Sánchez1, Beatriz Morillo Gutiérrez1, Belén Mosquera Pérez1, Pablo Obando Pacheco1, Jacobo Pardo-Seco1,2, Sara Pischedda1,2, Irene RiveroCalle1, Carmen Rodríguez-Tenreiro1, Lorenzo Redondo-Collazo1, Antonio Salas Ellacuriaga1,2, Sonia Serén Fernández1, María del Sol Porto Silva1, Ana Vega1,3, Lucía Vilanova Trillo1.

1 Translational Pediatrics and Infectious Diseases, Pediatrics Department, Hospital Clínico Universitario de Santiago, Santiago de Compostela, Spain, and GENVIP Research Group (http://www.genvip.org), Instituto de Investigación Sanitaria de Santiago, Galicia, Spain.

2 Unidade de Xenética, Departamento de Anatomía Patolóxica e Ciencias Forenses, Instituto de Ciencias Forenses, Facultade de Medicina, Universidade de Santiago de Compostela, and GenPop Research Group, Instituto de Investigaciones Sanitarias (IDIS), Hospital Clínico Universitario de Santiago, Galicia, Spain

3 Fundación Pública Galega de Medicina Xenómica, Servizo Galego de Saúde (SERGAS), Instituto de Investigaciones Sanitarias (IDIS), and Grupo de Medicina Xenómica, Centro de Investigación Biomédica en Red de Enfermedades Raras (CIBERER), Universidade de Santiago de Compostela (USC), Santiago de Compostela, Spain

#### EUCLIDS SPANISH CLINICAL NETWORK

Susana Beatriz Reyes1, María Cruz León León1, Álvaro Navarro Mingorance1, Xavier Gabaldó Barrios1, Eider Oñate Vergara2, Andrés Concha Torre3, Ana Vivanco3, Reyes Fernández3, Francisco Giménez Sánchez4, Miguel Sánchez Forte4, Pablo Rojo5, J.Ruiz Contreras5, Alba Palacios5, Cristina Epalza Ibarrondo5, Elizabeth Fernández Cooke5, Marisa Navarro6, Cristina Álvarez Álvarez6, María José Lozano6, Eduardo Carreras7, Sonia Brió Sanagustín7, Olaf Neth8, Mª del Carmen Martínez Padilla9, Luis Manuel Prieto Tato10, Sara Guillén10, Laura Fernández Silveira11, David Moreno12.

1 Hospital Clínico Universitario Virgen de la Arrixaca; Murcia, Spain.

2 Hospital de Donostia; San Sebastián, Spain.

3 Hospital Universitario Central de Asturias; Asturias, Spain.

4 Complejo Hospitalario Torrecárdenas; Almería, Spain.

5 Hospital Universitario 12 de Octubre; Madrid, Spain.

6 Hospital General Universitario Gregorio Marañón; Madrid, Spain.

7 Hospital de la Santa Creu i Sant Pau; Barcelona, Spain.

8 Hospital Universitario Virgen del Rocío; Sevilla, Spain.

9 Complejo Hospitalario de Jaén; Jaén, Spain.

10 Hospital Universitario de Getafe; Madrid, Spain.

11 Hospital Universitario y Politécnico de La Fe; Valencia, Spain.

12 Hospital Regional Universitario Carlos Haya; Málaga, Spain.

### Members of the Pediatric Dutch Bacterial Infection Genetics (PeD-BIG) network (the Netherlands)

#### Steering committee

Coordination: R. de Groot 1, A.M. Tutu van Furth 2, M. van der Flier 1

Coordination Intensive Care: N.P. Boeddha 3, G.J.A. Driessen 3, M. Emonts 3, 4, 5, J.A. Hazelzet 3

Other members: T.W. Kuijpers 7, D. Pajkrt 7, E.A.M. Sanders 6, D. van de Beek 8, A. van der Ende 8

Trial coordinator: H.L.A. Philipsen 1

### Local investigators (in alphabetical order)

A.O.A. Adeel 9, M.A. Breukels 10, D.M.C. Brinkman 11, C.C.M.M. de Korte 12, E. de Vries 13, W.J. de Waal 15, R. Dekkers 15, A. Dings-Lammertink 16, R.A. Doedens 17, A.E. Donker 18, M. Dousma19, T.E. Faber 20, G.P.J.M. Gerrits21, J.A.M. Gerver 22, J. Heidema 23, J. Homan-van der Veen 24, M.A.M. Jacobs 25, N.J.G. Jansen 6, P. Kawczynski 26, K. Klucovska 27, M.C.J. Kneyber 28, Y. Koopman-Keemink 29, V.J. Langenhorst 30, J. Leusink 31, B.F. Loza 32, I.T. Merth 33, C.J. Miedema 34, C. Neeleman 1, J.G. Noordzij 35, C.C. Obihara 36, A.L.T. van Overbeek – van Gils 37, G.H. Poortman 38,S.T. Potgieter 39, J. Potjewijd 40, P.P.R. Rosias 41, T. Sprong 21, G.W. ten Tussher 42, B.J. Thio 43, G.A. Tramper-Stranders 44, M. van Deuren 1, H. van der Meer 2, A.J.M. van Kuppevelt 45, A.M. van Wermeskerken 46, W.A. Verwijs 47, T.F.W. Wolfs 4.

1. Radboud University Medical Center – Amalia Children’s Hospital, Nijmegen, The Netherlands

2. Vrije Universiteit University Medical Center, Amsterdam, The Netherlands

3. Erasmus Medical Center – Sophia Children’s Hospital, Rotterdam, The Netherlands

4. Institute of Cellular Medicine, Newcastle University, Newcastle upon Tyne, United Kingdom

5. Paediatric Infectious Diseases and Immunology Department, Newcastle upon Tyne Hospitals Foundation Trust, Great North Children's Hospital, Newcastle upon Tyne, United Kingdom

6. University Medical Center Utrecht – Wilhelmina Children’s Hospital, Utrecht, The Netherlands

7. Academic Medical Center – Emma Children’s Hospital, University of Amsterdam, Amsterdam, The Netherlands

8. Academic Medical Center, University of Amsterdam, Amsterdam, The Netherlands

9. Kennemer Gasthuis, Haarlem, The Netherlands

10. Elkerliek Hospital, Helmond, The Netherlands

11. Alrijne Hospital, Leiderdorp, The Netherlands

12. Beatrix Hospital, Gorinchem, The Netherlands

13. Jeroen Bosch Hospital, ‘s-Hertogenbosch, The Netherlands

14. Diakonessenhuis, Utrecht, The Netherlands

15. Maasziekenhuis Pantein, Boxmeer, The Netherlands

16. Gelre Hospitals, Zutphen, The Netherlands

17. Martini Hospital, Groningen, The Netherlands

18. Maxima Medical Center, Veldhoven, The Netherlands

19. Gemini Hospital, Den Helder, The Netherlands

20. Medical Center Leeuwarden, Leeuwarden, The Netherlands

21. Canisius-Wilhelmina Hospital, Nijmegen, The Netherlands

22. Rode Kruis Hospital, Beverwijk, The Netherlands

23. St. Antonius Hospital, Nieuwegein, The Netherlands

24. Deventer Hospital, Deventer, The Netherlands

25. Slingeland Hospital, Doetinchem, The Netherlands

26. Refaja Hospital, Stadskanaal, The Netherlands

27. Bethesda Hospital, Hoogeveen, The Netherlands

28. University Medical Center Groningen, Beatrix Children’s hospital, Groningen, The Netherlands

29. Haga Hospital – Juliana Children’s Hospital, Den Haag, The Netherlands

30. Isala Hospital, Zwolle, The Netherlands

31. Bernhoven Hospital, Uden, The Netherlands

32. VieCuri Medical Center, Venlo, The Netherlands

33. Ziekenhuisgroep Twente, Almelo-Hengelo, The Netherlands

34. Catharina Hospital, Eindhoven, The Netherlands

35. Reinier de Graaf Gasthuis, Delft, The Netherlands

36. ETZ Elisabeth, Tilburg, The Netherlands

37. Scheper Hospital, Emmen, The Netherlands

38. St. Jansdal Hospital, Hardewijk, The Netherlands

39. Laurentius Hospital, Roermond, The Netherlands

40. Isala Diaconessenhuis, Meppel, The Netherlands

41. Zuyderland Medical Center, Sittard-Geleen, The Netherlands

42. Westfriesgasthuis, Hoorn, The Netherlands

43. Medisch Spectrum Twente, Enschede, The Netherlands

44. St. Franciscus Gasthuis, Rotterdam, The Netherlands 5

45. Streekziekenhuis Koningin Beatrix, Winterswijk, The Netherlands

46. Flevo Hospital, Almere, The Netherlands

47. Zuwe Hofpoort Hospital, Woerden, The Netherlands

### Swiss Pediatric Sepsis Study

#### Steering Committee

Luregn J Schlapbach, MD, FCICM 1,2,3, Philipp Agyeman, MD 1, Christoph Aebi, MD 1, Christoph Berger, MD 1 Luregn J Schlapbach, MD, FCICM 1,2,3, Philipp Agyeman, MD 1, Christoph Aebi, MD 1, Eric Giannoni, MD 4,5, Martin Stocker, MD 6, Klara M Posfay-Barbe, MD 7, Ulrich Heininger, MD 8, Sara Bernhard-Stirnemann, MD 9, Anita Niederer-Loher, MD 10, Christian Kahlert, MD 10, Paul Hasters, MD 11, Christa Relly, MD 12, Walter Baer, MD 13, Christoph Berger, MD 12 **for the Swiss Pediatric Sepsis Study**

1. Department of Pediatrics, Inselspital, Bern University Hospital, University of Bern, Switzerland

2. Paediatric Critical Care Research Group, Mater Research Institute, University of Queensland, Brisbane, Australia

3. Paediatric Intensive Care Unit, Lady Cilento Children’s Hospital, Children’s Health Queensland, Brisbane, Australia

4. Service of Neonatology, Lausanne University Hospital, Lausanne, Switzerland

5. Infectious Diseases Service, Lausanne University Hospital, Lausanne, Switzerland

6. Department of Pediatrics, Children’s Hospital Lucerne, Lucerne, Switzerland

7. Pediatric Infectious Diseases Unit, Children’s Hospital of Geneva, University Hospitals of Geneva, Geneva, Switzerland

8. Infectious Diseases and Vaccinology, University of Basel Children’s Hospital, Basel, Switzerland

9. Children’s Hospital Aarau, Aarau, Switzerland

10. Division of Infectious Diseases and Hospital Epidemiology, Children’s Hospital of Eastern Switzerland St. Gallen, St. Gallen, Switzerland

11. Department of Neonatology, University Hospital Zurich, Zurich, Switzerland

12. Division of Infectious Diseases and Hospital Epidemiology, and Children’s Research Center, University Children’s Hospital Zurich, Switzerland

13. Children’s Hospital Chur, Chur, Switzerland

### Liverpool Partner

#### Principal Investigators

Enitan Carrol1

Stéphane Paulus 1,2

ALDER HEY SERIOUS PAEDIATRIC INFECTION RESEARCH GROUP (ASPIRE) (in alphabetical order):

Hannah Frederick3, Rebecca Jennings3, Joanne Johnston3, Rhian Kenwright3

1 Department of Clinical Infection, Microbiology and Immunology, University of Liverpool Institute of Infection and Global Health, Liverpool, England

2 Alder Hey Children’s Hospital, Department of Infectious Diseases, Eaton Road, Liverpool, L12 2AP

3 Alder Hey Children’s Hospital, Clinical Research Business Unit, Eaton Road, Liverpool, L12 2AP

### Micropathology Ltd

Colin G Fink1,2, Elli Pinnock1

1 Micropathology Ltd Research and Diagnosis

2 University of Warwick

### Newcastle partner

Principle Investigator

Marieke Emonts1,2

Co-Investigator

Rachel Agbeko1,3

1 Institute of Cellular Medicine, Newcastle University, Newcastle upon Tyne, United Kingdom

2 Paediatric Infectious Diseases and Immunology Department, Newcastle upon Tyne Hospitals Foundation Trust, Great North Children's Hospital, Newcastle upon Tyne, United Kingdom

3 Paediatric Intensive Care Unit, Newcastle upon Tyne Hospitals Foundation Trust, Great North Children's Hospital, Newcastle upon Tyne, United Kingdom

### Gambia partner

Suzanne Anderson: Principal Investigator and West African study oversight:

Fatou Secka: Clinical research fellow and study co-ordinator

Additional Gambia site team (consortium members):

Kalifa Bojang: co-PI

Isatou Sarr: Senior laboratory technician

Ngange Kebbeh: Junior laboratory technician

Gibbi Sey: lead research nurse Medical Research Council Clinic

Momodou Saidykhan: lead research nurse Edward Francis Small Teaching Hospital

Fatoumata Cole: Data manager

Gilleh Thomas: Data manager

Martin Antonio: Local collaborator 7

### Austrian partner

#### PI

Werner Zenz1

Co-Investigators/Steering committee:

Daniela S. Klobassa1, Alexander Binder1, Nina A. Schweintzger1, Manfred Sagmeister1

1 University Clinic of Paediatrics and Adolescent Medicine, Department of General Paediatrics, Medical University Graz, Austria

### Austrian network, participating centres in Austria, Germany, Italy, Serbia, Lithuania, patient recruitment (in alphabetical order)

Hinrich Baumgart1, Markus Baumgartner2, Uta Behrends3, Ariane Biebl4, Robert Birnbacher5, Jan-Gerd Blanke6, Carsten Boelke7, Kai Breuling3, Jürgen Brunner8, Maria Buller9, Peter Dahlem10, Beate Dietrich11, Ernst Eber12, Johannes Elias13, Josef Emhofer2, Rosa Etschmaier14, Sebastian Farr15, Ylenia Girtler16, Irina Grigorow17, Konrad Heimann18, Ulrike Ihm19, Zdenek Jaros20, Hermann Kalhoff21, Wilhelm Kaulfersch22, Christoph Kemen23, Nina Klocker24, Bernhard Köster25, Benno Kohlmaier26, Eleni Komini27, Lydia Kramer3, Antje Neubert28, Daniel Ortner29, Lydia Pescollderungg16, Klaus Pfurtscheller30, Karl Reiter31, Goran Ristic32, Siegfried Rödl30, Andrea Sellner26, Astrid Sonnleitner26, Matthias Sperl33, Wolfgang Stelzl34, Holger Till1, Andreas Trobisch26, Anne Vierzig35, Ulrich Vogel12, Christina Weingarten36, Stefanie Welke37, Andreas Wimmer38, Uwe Wintergerst39, Daniel Wüller40, Andrew Zaunschirm41, Ieva Ziuraite42, Veslava Žukovskaja42

1 Department of Pediatric and Adolescence Surgery, Division of General Pediatric Surgery, Medical University Graz, Austria

2 Department of Pediatrics, General Hospital of Steyr, Austria

3 Department of Pediatrics/Department of Pediatric Surgery, Technische Universität München (TUM), Munich, Germany

4 Department of Pediatrics, Kepler University Clinic, Medical Faculty of the Johannes Kepler University, Linz, Austria

5 Department of Pediatrics and Adolesecent Medicine LKH Villach, Austria

6 Department of Pediatrics and Adolescent Medicine and Neonatology, Hospital Ludmillenstift, Meppen, Germany

7 Hospital for Children's and Youth Medicine, Oberschwabenklinik, Ravensburg, Germany

8 Department of Pediatrics, Medical University Innsbruck, Austria

9 Clinic for Paediatrics and Adolescents Medicine, Sana Hanse-Klinikum Wismar, Germany

10 Departement of Pediatrics, Medical Center Coburg, Germany

11 University Medicine Rostock, Department of Pediatrics (UKJ), Rostock, Germany

12 Department of Pulmonology, Medical University Graz, Austria

13 Institute for Hygiene and Microbiology, University of Würzburg, Germany

14 Clinical Institute of Medical and Chemical Laboratory Diagnostics, Medical University Graz, Austria

15 Department of Pediatric Orthopedics and Adult Foot and Ankle Surgery, Orthopedic Hospital Speising, Vienna, Austria

16 Department of Paediatrics, Regional Hospital Bolzano, Italy

17Department of Pediatrics and Adolescent Medicine, General Hospital Hochsteiermark/Leoben, Austria

18 Department of Neonatology and Paediatric Intensive Care, Children's University Hospital, RWTH Aachen, Germany

19 Paediatric Intensive Care Unit, Department of Paediatric Surgery, Donauspital Vienna, Austria

20 Department of Pediatrics, General Public Hospital, Zwettl, Austria

21 Pediatric Clinic Dortmund, Germany

22 Department of Pediatrics and Adolescent Medicine, Klinikum Klagenfurt am Wörthersee, Klagenfurt, Austria

23 Catholic Children’s Hospital Wilhelmstift, Department of Pediatrics, Hamburg, Germany 24 Department of Pediatrics, Krankenhaus Dornbirn, Austria

25 Children’s Hospital Luedenscheid, Maerkische Kliniken, Luedenscheid, Germany

26 Department of General Paediatrics, Medical University Graz, Austria

27 Department of Paediatrics, Schwarzwald-Baar-Hospital, Villingen-Schwenningen, Germany

28 Department of Paediatrics and Adolescents Medicine, University Hospital Erlangen, Germany

29 Department of Pediatrics and Adolescent Medicine, Medical University of Salzburg, Austria

30 Paediatric Intensive Care Unit, Medical University Graz, Austria

31 Dr. von Hauner Children's Hospital, Ludwig-Maximilians-Universitaet, Munich, Germany

32 Mother and Child Health Care Institute of Serbia, Belgrade, Serbia

33 Department of Pediatric and Adolescence Surgery, Division of Pediatric Orthopedics, Medical University Graz, Austria

34 Department of Pediatrics, Academic Teaching Hospital, Landeskrankenhaus Feldkirch, Austria

35 University Children’s Hospital, University of Cologne, Germany

36 Department of Pediatrics and Adolescent Medicine Wilheminenspital, Vienna, Austria

37 Department of Pediatric Surgery, Municipal Hospital Karlsruhe, Germany

38 Hospital of the Sisters of Mercy Ried, Department of Pediatrics and Adolescent Medicine, Ried, Austria

39 Hospital St. Josef, Braunau, Austria

40 Christophorus Kliniken Coesfeld Clinic for Pediatrics, Coesfeld, Germany

41 Department of Paediatrics, University Hospital Krems, Karl Landsteiner University of Health Sciences, Krems, Austria 42Children‘s Hospital, Affiliate of Vilnius University Hospital Santariskiu Klinikos, Lithuania

